# Genome-wide haplotype-based association analysis of major depressive disorder in Generation Scotland and UK Biobank

**DOI:** 10.1101/068643

**Authors:** David M. Howard, Lynsey S. Hall, Jonathan D. Hafferty, Yanni Zeng, Mark J. Adams, Toni-Kim Clarke, David J. Porteous, Reka Nagy, Caroline Hayward, Blair H. Smith, Alison D. Murray, Niamh M. Ryan, Kathryn L. Evans, Chris S. Haley, Ian J. Deary, Pippa A. Thomson, Andrew M. McIntosh

**Affiliations:** Division of Psychiatry, University of Edinburgh, Royal Edinburgh Hospital, Edinburgh, UK; Medical Research Council Human Genetics Unit, Institute of Genetics and Molecular Medicine, University of Edinburgh, Edinburgh, UK; Centre for Genomic and Experimental Medicine, Institute of Genetics and Molecular Medicine, University of Edinburgh, Edinburgh, UK; Division of Population Health Sciences, University of Dundee, Dundee, UK; Aberdeen Biomedical Imaging Centre, University of Aberdeen, Aberdeen, UK; Department of Psychology, The University of Edinburgh, Edinburgh, UK; Centre for Cognitive Ageing and Cognitive Epidemiology, The University of Edinburgh, Edinburgh, UK; Generation Scotland, Institute of Genetics and Molecular Medicine, University of Edinburgh, Edinburgh, UK

**Keywords:** Haplotype association analysis, Major Depressive Disorder, 6q21, Depression, Generation Scotland, UK Biobank

## Abstract

Genome-wide association studies using genotype data have had limited success in the identification of variants associated with major depressive disorder (MDD). Haplotype data provide an alternative method for detecting associations between variants in weak linkage disequilibrium with genotyped variants and a given trait of interest. A genome-wide haplotype association study for MDD was undertaken utilising a family-based population cohort, Generation Scotland: Scottish Family Health Study (n = 18 773), as a discovery cohort with UK Biobank used as a population-based cohort replication cohort (n = 25 035). Fine mapping of haplotype boundaries was used to account for overlapping haplotypes potentially tagging the same causal variant. Within the discovery cohort, two haplotypes exceeded genome-wide significance (P < 5 × 10^-8^) for an association with MDD. One of these haplotypes was nominally significant in the replication cohort (P < 0.05) and was located in 6q21, a region which has been previously associated with bipolar disorder, a psychiatric disorder that is phenotypically and genetically correlated with MDD. Several haplotypes with P < 10^^-7^^ in the discovery cohort were located within gene coding regions associated with diseases that are comorbid with MDD. Using such haplotypes to highlight regions for sequencing may lead to the identification of the underlying causal variants.

## INTRODUCTION

Major depressive disorder (MDD) is a complex and clinically heterogeneous condition with core symptoms of low mood and/or anhedonia over a period of at least two weeks. MDD is frequently comorbid with other clinical conditions, such as cardiovascular disease,^1^ cancer^2^ and inflammatory diseases.^3^ This complexity and comorbidity suggests heterogeneity of aetiology and may explain why there has been limited success in identifying causal genetic variants,^4-7^ despite heritability estimates ranging from 28% to 37%.^8, 9^ Single nucleotide polymorphism (SNP)-based analyses are unlikely to fully capture the variation in regions surrounding the genotyped markers, including untyped lower-frequency variants and those that are in weak linkage disequilibrium (LD) with the common SNPs on many genotyping arrays.

Haplotype-based analysis may help improve the detection of causal genetic variants as, unlike single SNP-based analysis, it is possible to assign the strand of sequence variants and combine information from multiple SNPs to identify rarer causal variants. A number of studies^10-12^ have identified haplotypes associated with MDD, albeit by focussing on particular regions of interest. In the current study, a family and population-based cohort Generation Scotland: Scottish Family Health Study (GS:SFHS) was utilised to ascertain genome-wide haplotypes in closely and distantly related individuals.^13^ A haplotype-based association analysis was conducted using MDD as a phenotype, followed by additional fine-mapping of haplotype boundaries with a replication and meta-analysis performed using the UK Biobank cohort.^14^

## MATERIALS AND METHODS

### Discovery cohort

The discovery phase of the study used the family and population-based Generation Scotland: Scottish Family Health Study (GS:SFHS) cohort,^13^ consisting of 23 960 individuals of whom 20 195 were genotyped with the Illumina OmniExpress BeadChip (706 786 SNPs). Individuals with a genotype call rate < 98% were removed, as well as those SNPs with a call rate < 98%, a minor allele frequency (MAF) < 0.01 or those deviating from Hardy-Weinberg equilibrium (*P* < 10^−6^). Individuals who were identified as population outliers through principal component analyses of their genotypic information were also removed.^15^

Following quality control there were 19 904 GS:SFHS individuals (11 731 females and 8 173 males) that had genotypic information for 561 125 autosomal SNPs. These individuals ranged from 18-99 years of age with an average age of 47.4 years and a standard deviation of 15.0 years. There were 4 933 families that had at least two related individuals, this included 1 799 families with two members, 1 216 families with three members and 829 families with four members. The largest family group consisted of 31 related individuals and there were 1 789 individuals that had no other family members within GS:SFHS.

### Replication cohort

The population-based UK Biobank^16^ (provided as part of project #4844) was used as a replication cohort to assess those haplotypes within GS:SFHS with *P* < 10^−6^. The UK Biobank data consisted of 152 249 individuals with genomic data for 72 355 667 imputed variants.^17^ The SNPs genotyped in GS:SFHS were extracted from the UK Biobank data and those variants with an imputation infoscore < 0.8 were removed, leaving 555 782 variants in common between the two cohorts. Those genotyped individuals listed as non-white British and those that had also participated in GS:SFHS were removed from within UK Biobank, leaving a total of 119 955 individuals.

### Genotype phasing and haplotype formation

The genotype data for GS:SFHS and UK Biobank was phased using SHAPEIT v2.r837.^18^ Genome-wide phasing was conducted on the GS:SFHS cohort, whilst the phasing of UK Biobank was conducted on a 50Mb window centred on those haplotypes identified within GS:SFHS with P < 10^−6^. The relatedness within GS:SFHS made it suitable for the application of the duoHMM method, which improves phasing accuracy by also incorporating family information.^19^ The default window size of 2Mb was used for UK Biobank and a 5Mb window was used for GS:SFHS as larger window sizes have been demonstrated to be beneficial when there is increased identity by descent (IBD) in the population.^18^ The number of conditioning states per SNP was increased from the default of 100 states to 200 states to improve phasing accuracy, with the default effective population size of 15 000 used. To calculate the recombination rates between SNPs during phasing the HapMap phase II b37^20^ was used. This build was also used to partition the phased data into haplotypes.

Three window sizes (1cM, 0.5cM and 0.25cM) were used to establish the SNPs that formed each haplotype.^21^ Each window was then moved along the genome by a quarter of the respective window size. There were a total of 97 333 windows with a mean number of SNPs per window of 157, 79 and 34 for the 1cM, 0.5cM and 0.25cM windows, respectively. Windows that were less 5 SNPs in length were removed. Within each window, those haplotypes that had a minor allele frequency < 0.005 or that deviating from Hardy-Weinberg equilibrium (*P* < 10^−6^) were not tested for association. However, they were included within the alternative haplotype when assessing the remaining 2 618 094 haplotypes. The reported haplotype positions relate to the outermost SNPs within each haplotype are in base pair (bp) position according to GRCh37.

To approximate the number of independently segregating haplotypes the clump command within Plink v1.90 ^22^ was applied. This provides an estimation of the Bonferroni correction required for multiple testing. When applying an LD *r*^*2*^ threshold of < 0.4 there were 1 070 216 independently segregating haplotypes within GS:SFHS, equating to a *P*-value < 5 × 10^−8^ for genome-wide significance. This threshold is also frequently applied to SNP-based and sequence-based association studies to account for multiple testing.^23^

### Phenotype ascertainment and patient linkage

#### Discovery cohort

Within GS:SFHS a diagnosis of MDD was made using initial screening questions and the Structured Clinical Interview for the Diagnostic and Statistical Manual of Mental Disorders (SCID).^24^ The SCID is an internationally validated approach to identifying episodes of depression and was conducted by clinical nurses trained in its administration. Further details regarding this diagnostic assessment have been described previously.^25^ In this study, MDD was defined by at least one instance of a major depressive episode which initially identified 2 659 cases, 17 237 controls and 98 missing (phenotype unknown) individuals.

In addition, the psychiatric history of cases and controls was examined using record linkage to the Scottish Morbidity Record.^26^ Within the control group, 1 072 participants were found to have attended at least one psychiatry outpatient clinic and were excluded from the study. In addition, 47 of the MDD cases were found to have additional diagnoses of either bipolar disorder or schizophrenia in psychiatric inpatient records and were also excluded from the study. These participants had given prior consent for anonymised record linkage to routine administrative clinical data.

In total there were 2 605 MDD cases and 16 168 controls following the removal of individuals based on patient records and population stratification, equating to a prevalence of 13.9% for MDD in this cohort.

#### Replication Cohort

Within the UK Biobank cohort, 25 035 participants completed a touchscreen assessment of depressive symptoms and previous treatment. On the basis of their responses, diagnostic status was defined as either ‘probable single lifetime episode of major depression’ or ‘probable recurrent major depression (moderate and severe)’ and with control status defined as ‘no mood disorder’. In total there were 8 508 cases and 16 527 controls, equating to a trait prevalence of 34.0% in this cohort, after the removal of individuals with insufficient information or ambiguous phenotypes.^14^

### Statistical approach

#### Discovery cohort

A mixed linear model was used to conduct an association analysis using GCTA v1.25.0 ^27^:

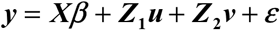

where y was the vector of binary observations for MDD.***β*** was the matrix of fixed effects, including haplotype, sex, age and age^2^. ***u*** was fitted as a random effect taking into account the genomic relationships (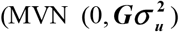, where ***G*** was a SNP-based genomic relationship matrix^28^). ***v*** was a random effect fitting a second genomic relationship matrix 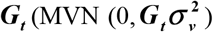 which modelled only the more closely related individuals.^29^ ***G***_*t*_ was equal to ***G*** except that off-diagonal elements < 0.05 were set to 0. ***X, Z***_1_ and ***Z***_2_ were the corresponding incidence matrices. ***ε*** was the vector of residual effects and was assumed to be normally distributed, 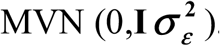.

The inclusion of the second genomic relationship matrix, ***G***_*t*_, was deemed desirable as the fitting of the single matrix G alone resulted in significant population stratification (intercept = 1.029 ± 0.003, λGC = 1.026) following examination with LD score regression.^30^ The fitting of both genomic relationship matrices simultaneously produced no evidence of bias due to population stratification (intercept = 1.002 ± 0.003, λGC = 1.005).

#### Replication cohort

A mixed linear model was used to assess the haplotypes in UK Biobank which were identified in the discovery cohort with *P* < 10^−6^ using GCTA v1.25.0 ^27^:

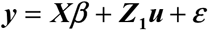

where***y*** was the vector of binary observations for MDD. ***β*** was the matrix of fixed effects, including haplotype, sex, age, age^2^, genotyping batch and recruitment centre. ***u*** was fitted as a random effect taking into account the SNP-based genomic relationships 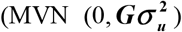. ***X*** and ***Z***_1_ were the corresponding incidence matrices and *ε* was the vector of residual effects and was assumed to be normally distributed, 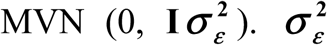. Replication success was judged on the statistical significance of each haplotype using an inverse variance-weighted meta-analysis across both cohorts conducted using Metal.^31^

#### Fine mapping

The method described above examines the effect of each haplotype against all other haplotypes in that window. Therefore, a haplotype could be assessed against similar haplotypes containing the same causal variant, limiting any observed phenotypic association. To investigate whether there were causal variants located within directly overlapping haplotypes of the same window size, fine mapping of haplotype boundaries was used. Where there were directly overlapping haplotypes, each with *P* < 10^−3^ and with an effect in the same direction, i.e. both causal or both preventative, then any shared consecutive regions formed a new haplotype that was assessed using the mixed model described previously. This new haplotype was assessed using all individuals and was required to be at least 5 SNPs in length. A total of 47 new haplotypes were assessed from within 26 pairs of directly overlapping haplotypes.

## RESULTS

An association analysis for MDD was conducted using 2 618 094 haplotypes and 47 fine mapped haplotypes within the discovery cohort, GS:SFHS. A genome-wide Manhattan plot of –log__10__ *P*-values for these haplotypes is provided in Figure 1 with a q-q plot provided in Supplementary Figure S1. Within the discovery cohort, two haplotypes exceeded genome-wide significance (*P* < 5 × 10^−8^) for an association with MDD, one located on chromosome 6 and the other located on chromosome 10. There were 12 haplotypes with *P* < 10^−6^ in the discovery cohort with replication sought for these haplotypes using UK Biobank. Summary statistics from both cohorts and the meta-analysis for these 12 haplotypes are provided in Table 1. The protein coding genes which overlap these 12 haplotypes along with the observed haplotype frequencies within the two cohorts are provided in Table 2. The SNPs and alleles that constitute these 12 haplotypes are provided in Supplementary Table S1.

**Figure 1.**
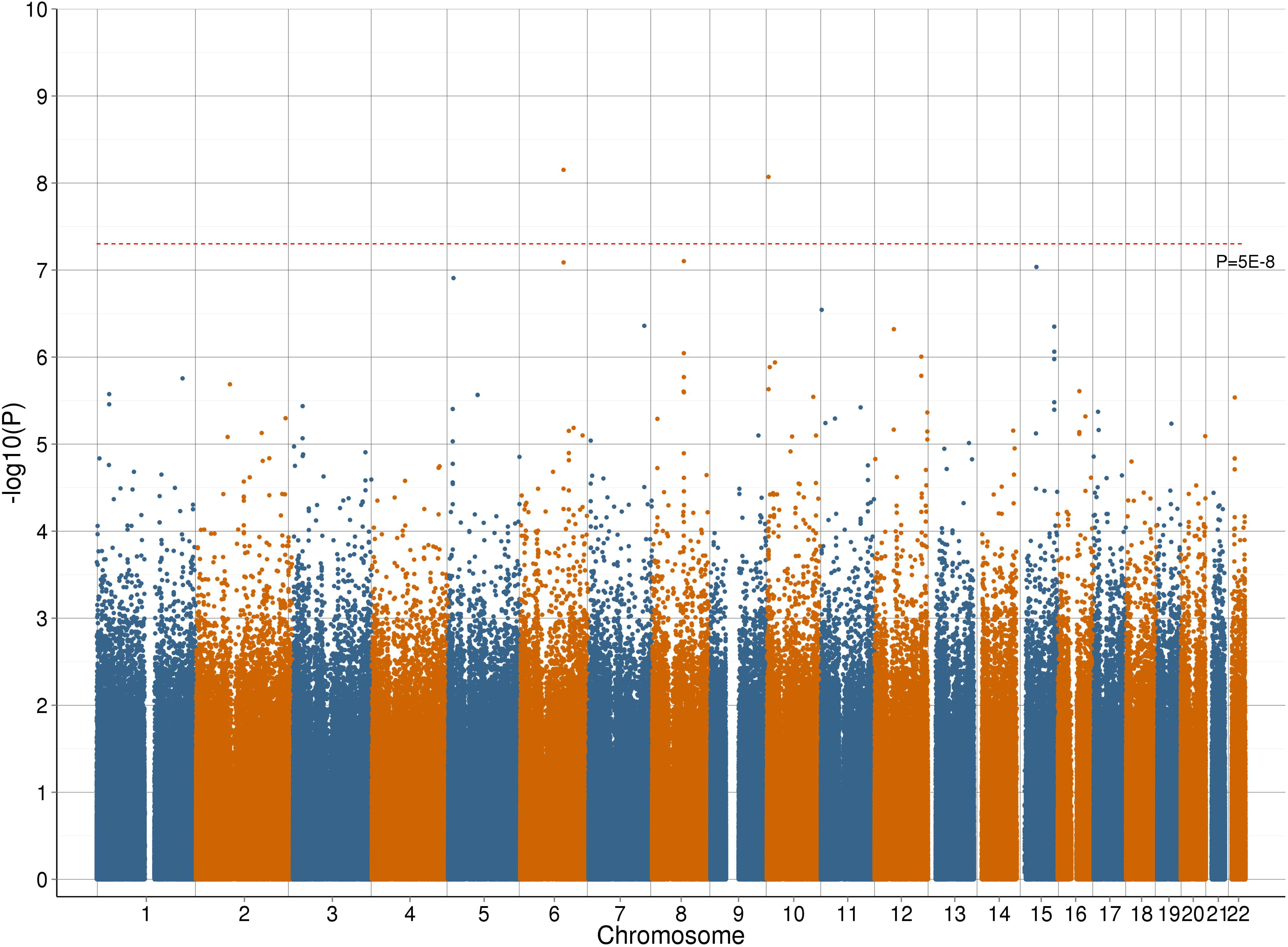
Manhattan plot representing the –log__10__ *P*-values for an association between each assessed haplotype in the Generation Scotland: Scottish Family Health Study cohort and Major Depressive Disorder

**Table 1.**
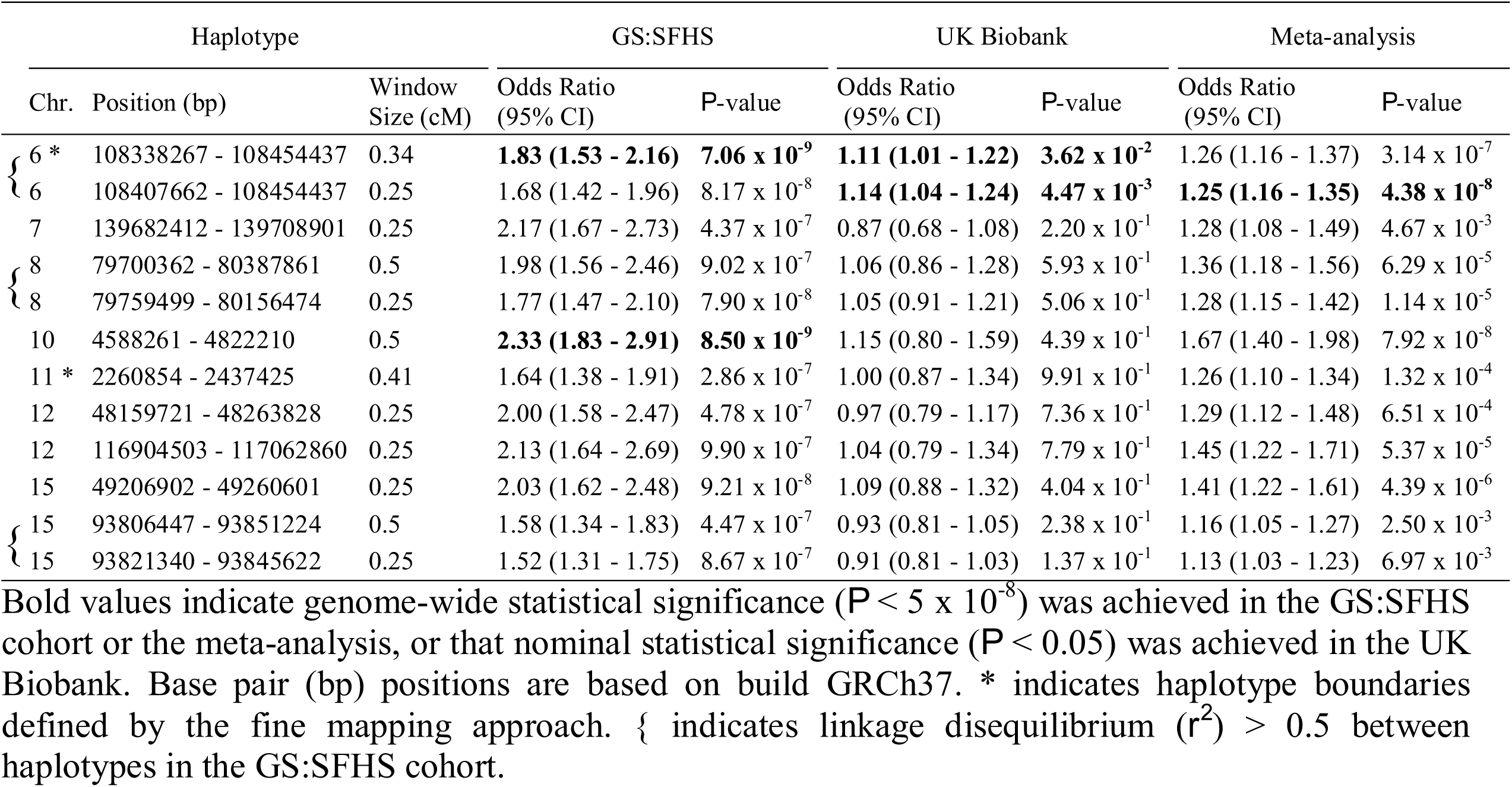
The genetic association between Major Depressive Disorder and 12 haplotypes in the Generation Scotland: Scottish Family Health Study (GS:SFHS) discovery cohort (where *P* < 10^−6^), the replication cohort (UK Biobank) and a meta-analysis.

**Table 2.**
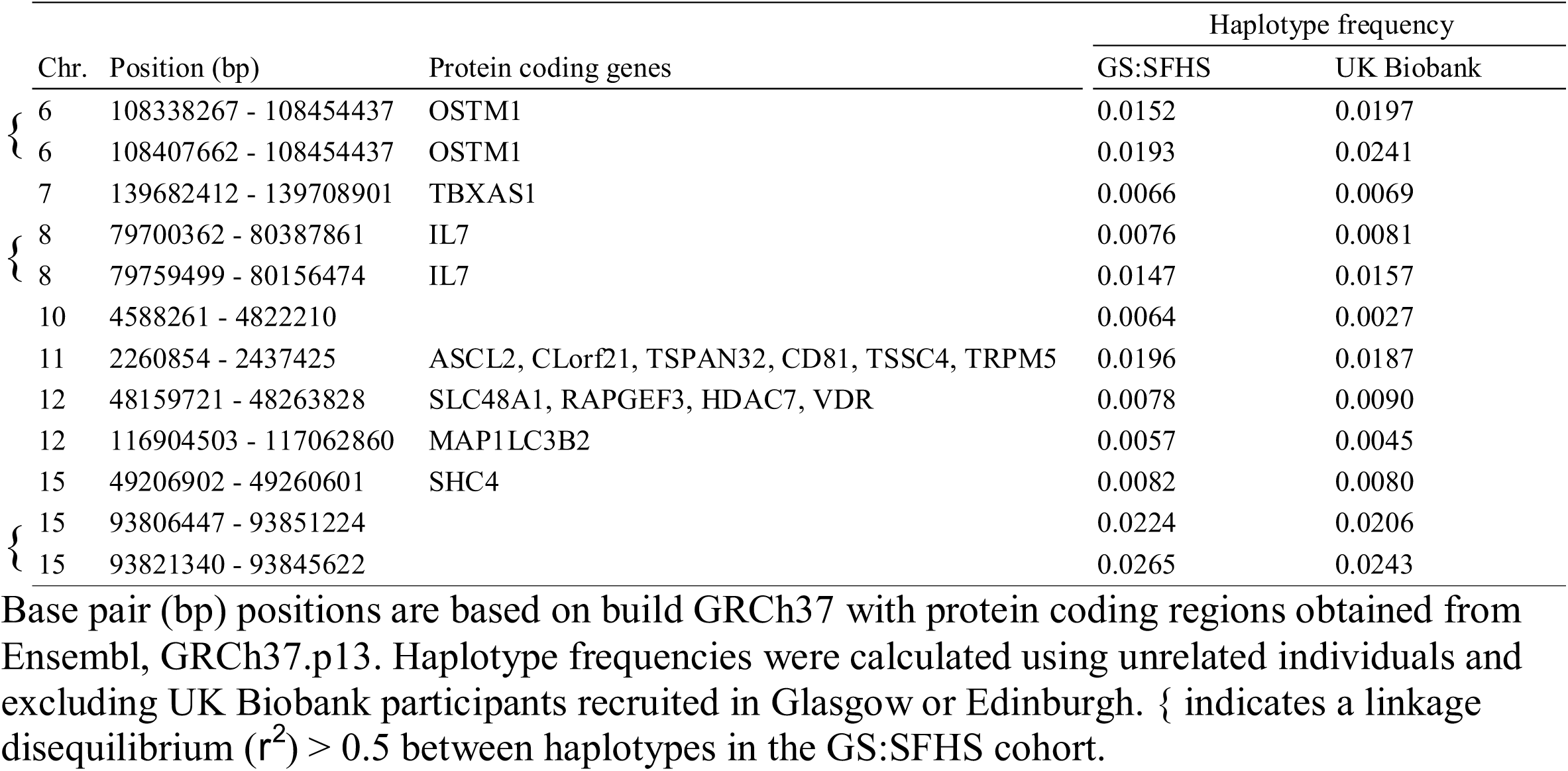
Protein coding genes located overlapping with the 12 haplotypes with *P* < 10^−6^ in the Generation Scotland: Scottish Family Health Study (GS:SFHS) discovery cohort and the frequencies of those haplotypes in GS:SFHS and UK Biobank.

The two haplotypes on chromosome 6 (LD *r*^*2*^ = 0.74) with *P* < 10^−6^ in the discovery cohort both achieved nominal significance (*P* < 0.05) in the replication cohort, with one reaching genome-wide significance (*P* < 5 × 10^−8^) in the meta-analysis. A regional association plot of the region surrounding these haplotypes within GS:SFHS is provided in Figure 2. Fine mapping was used to form the most significant haplotype within the discovery cohort. Two directly overlapping 0.5cM haplotypes consisting of 28 SNPs were identified between 108 335 345 and 108 454 437 bp (rs7749081 -rs212829). These two haplotypes had *P*-values of 3.24 × 10^−5^ and 5.57 × 10^−5^, respectively and differed at a single SNP (rs7749081). Exclusion of this single SNP defined a new 27 SNP haplotype that had a genome-wide significant association with MDD (*P* = 7.06 × 10^−9^). Calculating the effect size at the population level,^32^ the estimates of the contribution of the two haplotypes to the total genetic variance was 2.09 × 10^−4^ and 2.38 × 10^−4^, respectively, within GS:SFHS. None of the individual SNPs located within either haplotype were associated with MDD in either cohort (*P* ≥ 0.05).

**Figure 2.**
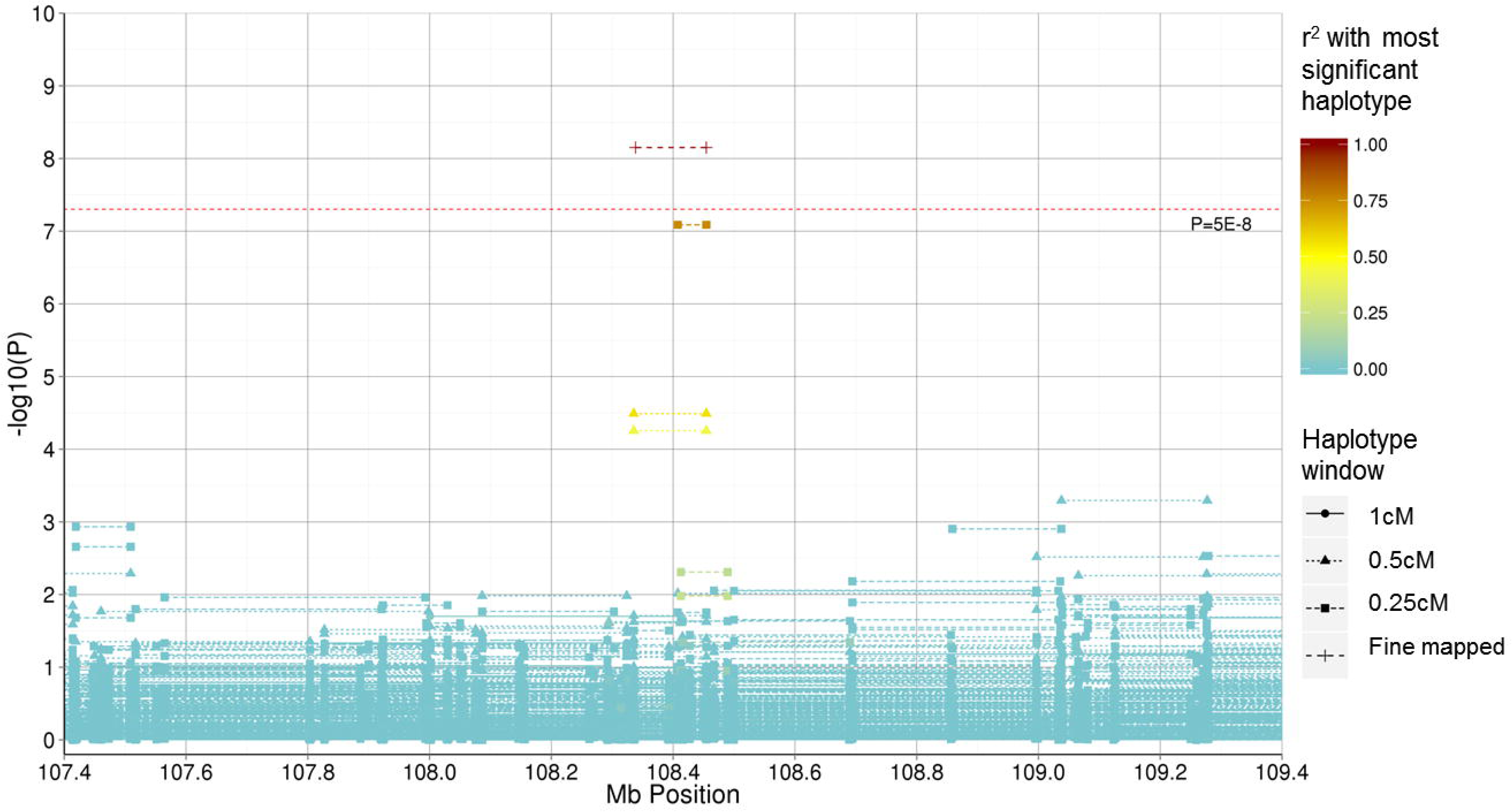
Regional association plot representing the –log__10__ *P*-values for an association between haplotypes in the Generation Scotland: Scottish Family Health Study cohort and Major Depressive Disorder within the 107.4 – 107.6 Mb region on chromosome 6. The start and end position (using build GRCh37) of haplotypes represent the outermost SNP positions within the windows examined. The warmth of colour represents the *r*^*2*^ with the genome-wide significant haplotype located between 108 338 267 and 108 454 437 bp.

A genome-wide significant haplotype (*P* = 8.50 × 10^−9^) was identified on chromosome 10 within GS:SFHS using a 0.5cM window. A regional association plot of the region surrounding this haplotype is provided in Figure 3. This haplotype had an odds ratio (OR) of 2.33 (95% CI: 1.83 – 2.91) in the discovery cohort and an OR of 1.15 (95% CI: 0.80 - 1.59) in the replication cohort. These were the highest ORs observed in the respective cohorts. The estimate of the contribution of this haplotype to the total genetic variance was 2.29 × 10^−4^ in the discovery cohort. Association analysis of the 92 SNPs on this haplotype revealed that one SNP in GS:SFHS (rs17133585) and two SNPs in UK Biobank (rs12413638 and rs10904290) were nominally significant (*P* < 0.05), although none had *P*-values < 0.001.

**Figure 3.**
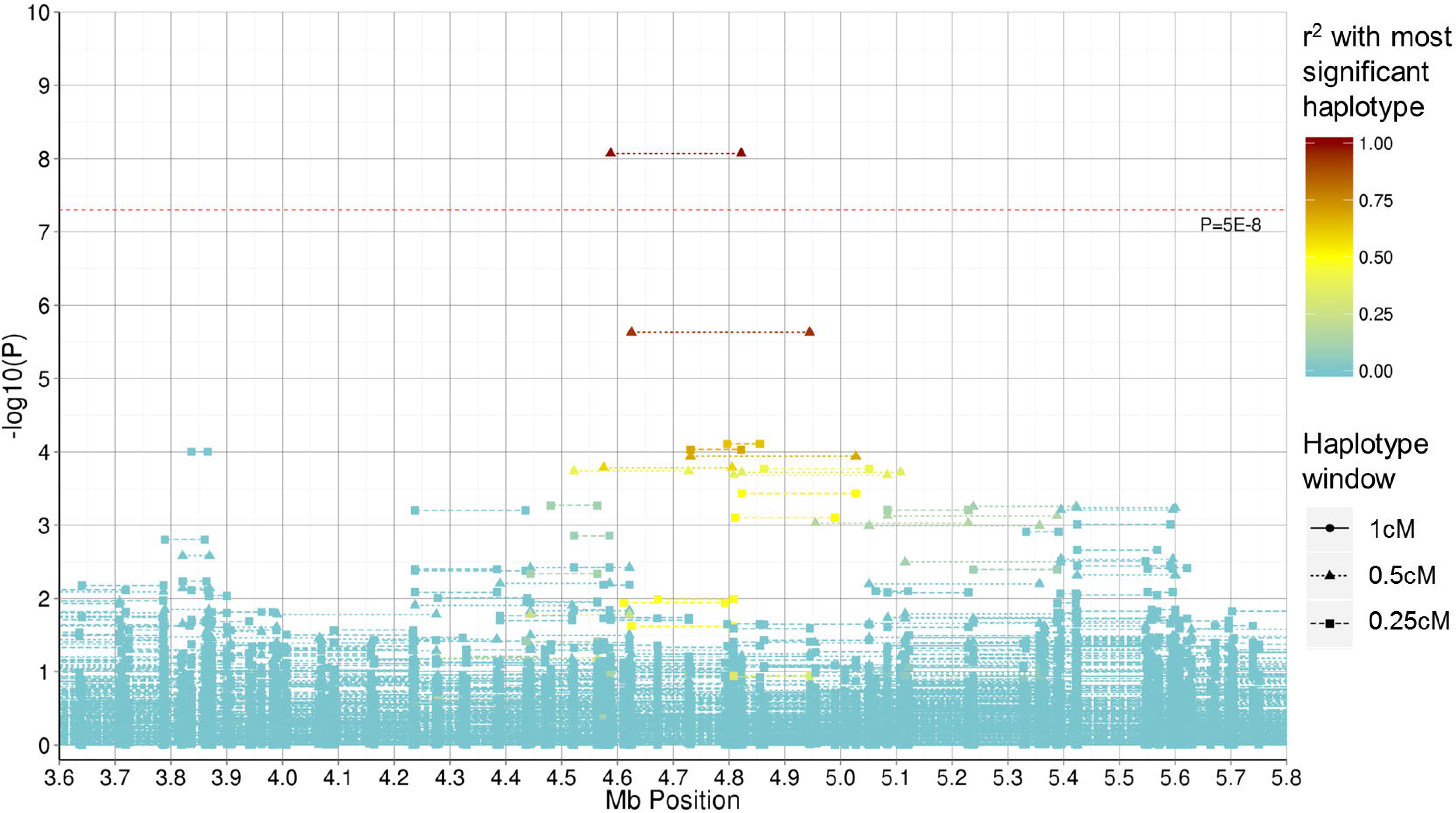
Region association plot representing the –log__10__ *P*-values for an association between haplotypes in the Generation Scotland: Scottish Family Health Study cohort and Major Depressive Disorder within the 3.6 – 5.8 Mb region on chromosome 10. The start and end position (using build GRCh37) of haplotypes represent the outermost SNP positions within the windows examined. The warmth of colour represents the *r*^*2*^ with the genome-wide significant haplotype located between 4 588 261 and 4 822 210 bp.

All 12 of the haplotypes with a *P*-value for association < 10^−6^ in the GS:SFHS discovery cohort were risk factors for MDD (OR > 1) and within the replication cohort, 7 out of these 12 haplotypes had OR > 1. None of the 95% confidence intervals for the replication ORs overlapped the 95% confidence intervals of the discovery GS:SFHS cohort.

## DISCUSSION

Twelve haplotypes were identified in the discovery cohort with *P* < 10^−6^ of which two were significant at the genome-wide level (*P* < 5 × 10^−8^) in the discovery cohort and one which was genome-wide significant (*P* < 5 × 10^−8^) in the meta-analysis. A power analysis^33^ was conducted using the genotype relative risks observed in the discovery cohort, the sample sizes and haplotype frequencies in the replication cohort and the prevalence of MDD reported for a structured clinical diagnosis of MDD in other high income counties (14.6%).^34^ There was sufficient power (> 0.99) to detect the twelve haplotypes with *P* < 10^−6^ identified in the discovery cohort within the replication cohort at a significance threshold of 0.05%.

A complementary approach to replication is to identify the gene coding regions within haplotypes that potentially provide a biologically informative explanation for an association with MDD. Those haplotypes with *P* < 10^−7^ in the discovery cohort and the gene coding regions that they overlap are discussed below.

The two haplotypes on chromosome 6 overlapped with the Osteopetrosis Associated Transmembrane Protein 1 (*OSTM1*) coding gene. *OSTM1* is associated with neurodegeneration^35, 36^ and melanocyte function,^37^ and alpha-melanocyte stimulating hormone has been shown to have an effect on depression-like symptoms.^38-40^ This haplotype lies within the 6q21 region that has been associated with bipolar disorder,^41-45^ a disease that shares symptoms with MDD and has a correlated phenotypic liability of 0.64.^46^ This may indicate either a pleiotropic effect or clinical heterogeneity, whereby patients may be misdiagnosed, i.e. patients may have MDD and transition to bipolar disorder in the future or are sub-threshold for bipolar disorder and instead given a diagnosis of MDD.

The haplotype identified on chromosome 8 overlapped with the Interleukin 7 (*IL7*) protein coding region. *IL7* is involved in maintaining T cell homeostasis^47^ and proliferation,^48^ which in turn contributes to the immune response to pathogens. It has been proposed that impaired T cell function may be a factor in the development of MDD,^49^ with depressed subjects found to have elevated^50^ or depressed levels^51^ of *IL7* serum. There is conjecture as to whether MDD causes inflammation or represents a reaction to an increased inflammatory response,^52, 53^ but it is most likely to be a bidirectional relationship.^51^

The haplotype on chromosome 10 overlapped with two RNA genes: long intergenic non-protein coding RNA 704 (*LINC00704*) and long intergenic non-protein coding RNA 705 (*LINC00705*). The function of these non-protein coding genes is unreported. However, a study of cardiac neonatal lupus which is a rare autoimmune disease demonstrated an association for a SNP (rs1391511) which is 15kb from *LINC00705*.

Two Dutch studies^54, 55^ have identified a variant (rs8023445) on chromosome 15 located within the SRC (Src homology 2 domain containing) family, member 4 (*SHC4*) gene coding region that has a moderate degree of association with MDD (*P* = 1.64 × 10^−5^ and *P* = 9 × 10^−6^, respectively). A variant (rs10519201) within the *SHC4* coding region was also found to have an association (*P* = 6.16 × 10^−6^) with Obsessive-Compulsive Personality Disorder in a UK-based study.^56^ *SHC4* is expressed in neurons^57^ and regulates BDNF-induced MAPK activation,^58^ which has been shown to be a key factor in MDD pathophysiology.^59^ The *SHC4* region overlaps with the haplotype on chromosome 15 identified in the discovery cohort (located at 49 206 902 – 49 260 601 bp) and therefore further research to examine the association between the *SHC4* region and psychiatric disorders could be warranted.

Haplotype-based analyses are capable of tagging variants due to the LD between the untyped variants and the multiple flanking genotyped variants which make up the inherited haplotype. This approach should provide greater power when there is comparatively higher IBD sharing, such as in GS:SFHS, where there is a greater likelihood that a single haplotype is tagging the same causal variant across that population. The UK Biobank was selected as replication cohort as it is a large population-based sample that was expected to be genetically similar to the GS:SFHS discovery cohort. This was confirmed by the similarity of the observed haplotype frequencies (Table 2) between the two cohorts. The prevalence of MDD observed in the discovery cohort (13.7%) was comparable to that reported (14.6%) within similar populations.^34^ However, in the replication cohort, the trait prevalence was notably higher (34.0%), most likely due to the differing methods of phenotypic ascertainment. Additional work could seek to replicate the findings in further cohorts, as well as full meta-analysis of all haplotypes within those cohorts. An additive model was used to analyse the haplotypes and alternative approaches could implement a dominant model or an analysis of diplotypes (haplotype pairs) for association with MDD.

## Conclusions

This study identified two haplotypes within the discovery cohort that exceeded genome-wide significance for association with a clinically diagnosed MDD phenotype. One of these haplotypes was nominally significant in the replication cohort and was in LD with a haplotype that was genome-wide significant in the meta-analysis. The genome-wide significant haplotype on chromosome 6 was located on 6q21, which has been shown previously to be related to psychiatric disorders. There were a number of haplotypes approaching genome-wide significance located within genic regions associated with diseases that are comorbid with MDD and therefore these regions warrant further investigation. The total genetic variance explained by the haplotypes identified was small, however these haplotypes potentially represent biologically informative aetiological subtypes for MDD and merit further analysis.

## ACKNOWLEDGEMENTS

Generation Scotland received core funding from the Chief Scientist Office of the Scottish Government Health Directorate CZD/16/6 and the Scottish Funding Council HR03006. Genotyping of GS:SFHS was carried out by the Genetics Core Laboratory at the Wellcome Trust Clinical Research Facility, Edinburgh, Scotland and was funded by the UK’s Medical Research Council and the Wellcome Trust (Wellcome Trust Strategic Award “Stratifying Resilience and Depression Longitudinally” (STRADL) (Reference 104036/Z/14/Z).

We are grateful to all the families who took part, the general practitioners and the Scottish School of Primary Care for their help in recruiting them, and the whole Generation Scotland team, which includes interviewers, computer and laboratory technicians, clerical workers, research scientists, volunteers, managers, receptionists, healthcare assistants and nurses. Ethics approval for the study was given by the NHS Tayside committee on research ethics (reference 05/S1401/8)

This research has been conducted using the UK Biobank resource – application number 4844; we are grateful to UK Biobank participants. The UK Biobank study was conducted under generic approval from the NHS National Research Ethics Service (approval letter dated 17th June 2011, Ref 11/NW/0382).

YZ acknowledges support from China Scholarship Council. IJD is supported by the Centre for Cognitive Ageing and Cognitive Epidemiology which is funded by the Medical Research Council and the Biotechnology and Biological Sciences Research Council (MR/K026992/1). AMMcI and T-KC acknowledges support from the Dr Mortimer and Theresa Sackler Foundation.

